# Metabolic pathways fuelling Devil Facial Tumour Disease

**DOI:** 10.1101/2025.03.12.642198

**Authors:** Anne-Lise Gérard, Florence Pirard, Caitlin Vanbeek, Antoine M. Dujon, Aaron G. Schultz, Rodrigo Hamede, Hannah V. Siddle, Frédéric Thomas, Matthew McKenzie, Mark Ziemann, Beata Ujvari

## Abstract

Devil Facial Tumour Diseases (DFTD), threatening Tasmanian devils, consist of two distinct transmissible cancers, DFT1 and DFT2, with differing origins and geographic spread. We investigated the metabolic differences between DFT1 and DFT2, examining cell viability, metabolic outputs, and bulk gene expression. Using both DFT1 and DFT2 cell lines and biopsies, we found that glycolysis, oxidative phosphorylation, glutamate metabolism and fatty acid synthesis are all essential for the survival of both tumour types. However, DFT2 exhibited higher rates of glycolysis and lactate generation compared to DFT1. This coincided with elevated ATP production, cholesterol biosynthesis and ROS generation, as well as an increased reliance on fatty acid metabolism. Furthermore, DFT2 is less metabolically adaptable than DFT1, being unable to switch to oxidative phosphorylation as DFT1 can when required. These metabolic changes in DFT2, in conjunction with its higher growth rate, suggests a more aggressive cancer phenotype than DFT1. Our findings highlight distinct metabolic adaptations in DFT2 that may contribute to its competitive advantage.

## INTRODUCTION

In multicellular organisms, cell division is tightly regulated to maintain tissue integrity and function. Such regulatory mechanisms comprise, but are not limited to, cell cycle checkpoints [1], telomere length [2] and immunosurveillance [3]. When one or several of these safeguards are impaired, cells proliferate uncontrollably, which can lead to cancer. Traits tumour cells acquire on the path to becoming cancerous are known as the hallmarks of cancer [4, 5]. One important hallmark is the deregulation of cellular metabolism, as tumour cells need to generate the necessary resources (nucleotides, building blocks for membranes, electron acceptors, etc.) to sustain high cell division rates. Cancer cells also evolve in a challenging environment where oxygen and nutrient availability vary over space and time with tumour size and vascularisation. Hence these cells must rewire their metabolism to produce enough precursors and energy, while also maintaining redox homeostasis by detoxifying excess reactive oxygen species (ROS) produced by the mitochondria [6].

The most common metabolic alteration found in cancer cells is increased glucose uptake and fermentation of glucose to lactate, also known as the “Warburg effect” [7]. Whilst normal cells completely oxidise glucose through the tricarboxylic acid (TCA) cycle and the electron transport chain (ETC; i.e., oxidative phosphorylation or OXPHOS), cancer cells convert most glucose to lactate through aerobic glycolysis [7]. The Warburg effect allows cancer cells to produce energy in the form of adenosine triphosphate (ATP) along with reducing equivalents at a higher rate compared to OXPHOS. Moreover, aerobic glycolysis has been shown to: (i) promote tissue invasion and immune evasion, (ii) drive the biosynthesis of nucleotides, lipids, and proteins, and finally (iii) play a role in cell signalling through ROS and chromatin modulation[8]. Alternatively, glucose can enter the pentose phosphate pathway (PPP) and participate in nucleotide synthesis. The PPP is also a major source of nicotinamide adenine dinucleotide phosphate (NADPH), contributing to ROS detoxification [7].

Aside from glucose, other molecules fuel cancer metabolism with glutamine being the second most consumed nutrient by cancer cells [7]. Most glutamine is converted to glutamate which can then be further processed to amino acids, nucleotides, or a-ketoglutarate, respectively contributing to protein synthesis, nucleic acid production and fuelling the TCA cycle [7, 9, 10]. Furthermore, cancer cells harness lipid metabolism to obtain sufficient energy (ATP), building blocks for their membranes, and signalling molecules for proliferation, survival, invasion, and metastasis [11]. Precursors and products of the TCA cycle such as glucose, acetyl coenzyme A (acetyl-CoA) and citrate can be used to produce lipids including fatty acids and cholesterol [7, 9]. Both enhanced lipid synthesis and lipid breakdown have been linked to cancer development [7].

These metabolic alterations occur not only during cancer development but also throughout the metastatic cascade (i.e., when cells detach from the extracellular matrix, travel through the bloodstream, and establish in a new tissue) [6]. An even more radical environmental change occurs in transmissible cancers where cancer cells are passed on from one individual to another as allo- or xeno-grafts [12–14]. Only two studies have investigated metabolic changes in transmissible cancers, one in bivalve transmissible neoplasia (BTN) [15] that impacts a variety of shellfish species and the other in Devil Facial Tumour 1 (DFT1) [16] that is endangering Tasmanian devils [17, 18]. The study on BTN used transcriptomics to identify key cancer drivers in bivalve neoplastic cells and found many metabolic pathways (including TCA and glycolysis) to be altered in BTN cells compared to healthy haemolymph cells [15]. The study on DFT1 used a more focused, assay-based approach and found that DFT1 depends on carbohydrate metabolism to proliferate [16]. DFT1 has resulted in 60% of population loss [19] and resulted in the species being listed as endangered on the IUCN Red List.

Devil Facial Tumour Diseases (DFTD) are two transmissible facial cancers affecting the Tasmanian devil (*Sarcophilus harrisii*) [14]. The diseases are fatal and passed on through biting, which is frequent during social interactions [20]. DFTD consist of two distinct cancers that arose independently in separate founder animals: DFT1, estimated to have emerged in 1986 in Northeast Tasmania, and DFT2, estimated to have emerged in 2011 in the Southeast of the island [14, 21, 22]. Both cancers cause facial tumours that eventually lead to animals dying from starvation or organ failure due to metastatic disease within six to twelve months after the appearance of visible tumours [21, 23, 24]. Although both cancers cause similar clinical symptoms, a few notable differences between them exist that include (i) devil- and cell type-of-origin, (ii) geographical spread and (iii) *in vitro* growth dynamics. First, DFT1 originated from a myelinating Schwann cell of a female devil, while DFT2 originated from an immature or repair Schwann cell of a male devil [25, 26]. Interestingly, repair Schwann cells show increased motility and proliferation compared to differentiated Schwann cells [27], potentially translating to more aggressive cancers prone to metastasise. Second, DFT1 has propagated to most of Tasmania while DFT2 has a restricted geographic range, only recently affecting devils outside of the peninsula where it originated to co-occur with DFT1 [28]. This has led to three reported cases of co-infection with both DFT1 and DFT2 in devils, and competition between the two diseases for hosts has been hypothesised [29]. In a recent *in vitro* study, DFT1 cells were observed to grow slower in comparison to DFT2 cells, with DFT2 outcompeting DFT1 in co-cultures [30].

Recent work by Gérard et al.[30] hypothesised that differences in DFT1 and DFT2 growth dynamics could be explained by the distinct differentiation state of their Schwann cell-of-origin [26]. For instance, less differentiated repair Schwann cells dramatically increase glycolysis to assist in peripheral nerve repair [27]. This should translate into metabolic differences ultimately responsible for DFT1 and DFT2’s diverging *in vitro* growth dynamics and competitiveness.

Here, we investigate similarities and differences between DFT1 and DFT2 metabolism by growing cell lines in the presence of metabolic inhibitors and measuring cell viability. We also measured outputs of major metabolic pathways such as cholesterol content, lactate, ATP, and ROS production. To link our functional assays to gene expression data, we generated full transcriptomes for DFT1 and DFT2 cell lines and reanalysed previously published transcriptomes from tumour biopsies taken from wild infected animals [26]. We highlight distinct metabolic adaptations in DFT2 that may underpin its increased proliferative capacity and competitive advantage over DFT1.

Though rare, transmissible cancers offer real-world insight into cancer survival under stress as they exhibit extreme metabolic flexibility, mirroring human cancers during metastasis and therapy resistance. Understanding their metabolic vulnerabilities could position DFTD cells as natural models for testing inhibitors and identifying biomarkers for aggressive or treatment-resistant cancers. Investigating them can reveal how tumours evolve and adapt metabolically, providing a unique window into cancer biology.

## RESULTS

### Many enriched pathways are shared between DFT cell lines and biopsies

To understand the metabolic differences between DFT1 and DFT2, we used bulk RNA-sequencing of DFT1 and DFT2 cell cultures and tumour biopsies. According to multidimensional scaling (MDS), tumour biopsies clustered into distinct groups, suggesting clear differences between DFT1 and DFT2 (Fig 1A). DFT1 and DFT2 cell lines also separated into two distinct clusters on the first dimension of the MDS plots (Fig. 1B). When combining both datasets, cell lines clustered separately from tumour biopsies on the first dimension, and DFT1 cell lines grouped away from DFT2 cell lines on the second dimension (Fig. 1C). Interestingly, the biopsies did not cluster according to DFT on the second dimension, indicating less variability between DFT1 and DFT2 biopsies than between DFT1 and DFT2 cell line gene expression profiles. Thus, DFT biopsies were more similar in gene expression than DFT cell lines. Biopsy tumour purity was comparable to cell line tumour purity (78.77% and 79.34% respectively; Table S1), indicating minimal contamination with healthy host tissue.

**Figure 1.**
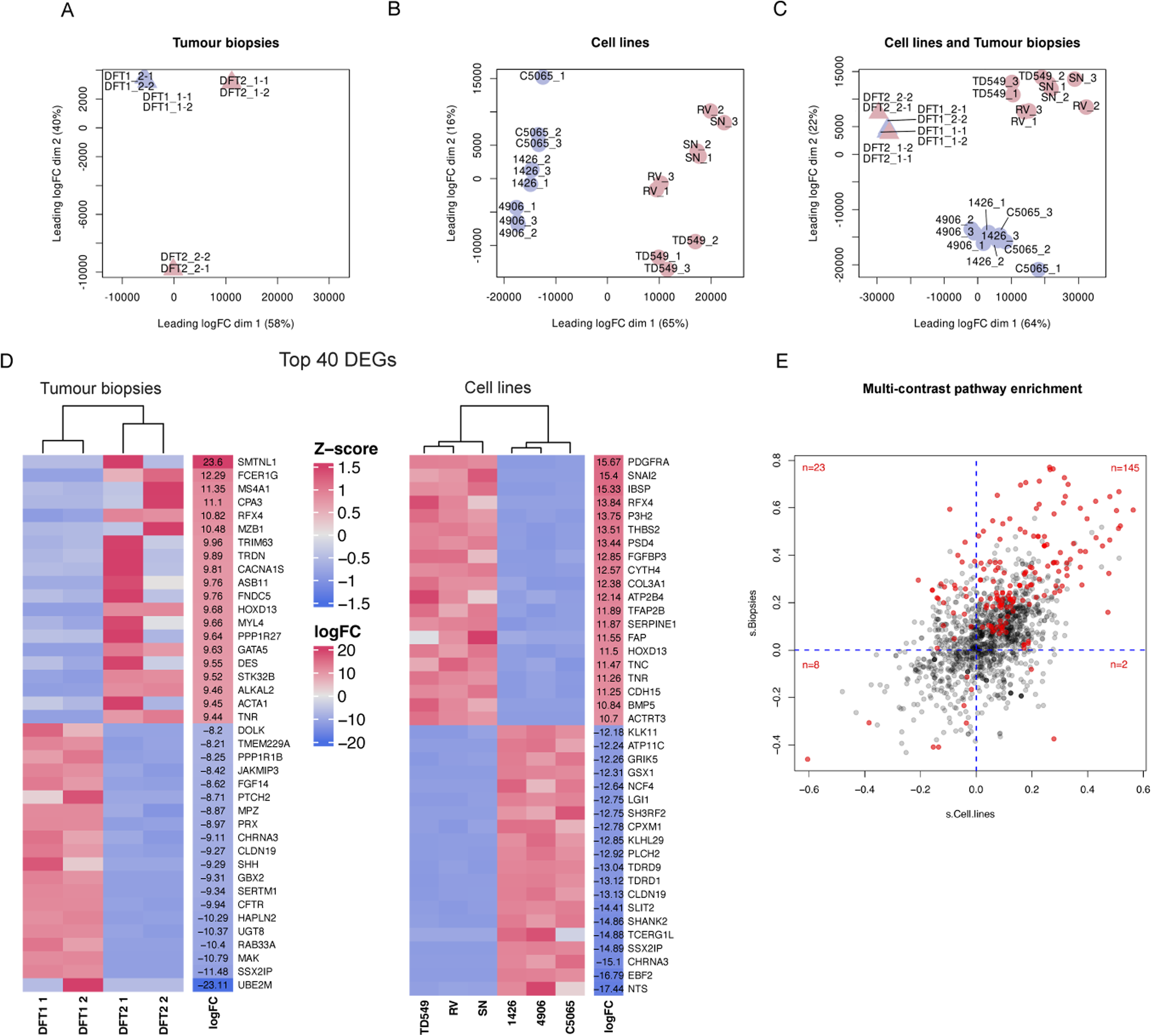
*In vitro* Devil Facial Tumour 1 (DFT1) and DFT2 transcriptomes differ more than *in vivo* ones. **A** Multi-dimensional scaling (MDS) plot of biopsy RNA sequences (Devil Facial Tumour 1, DFT1 in blue triangles and DFT2 in pink triangles). One experiment, 8 samples, 2 biological replicates per DFT, 2 technical replicates per biological replicate. **B** MDS plot of cell line RNA sequences (DFT1 in blue circles and DFT2 in pink circles). One experiment, 18 samples, 3 biological replicates per DFT, 3 technical replicates per biological replicate. **C** MDS plot of both cell line and biopsy RNA sequences. **D** Top 20 genes up- and top 20 genes down-regulated in DFT2, for biopsies and cell lines. **E** Multi-contrast enrichment analysis results. Each dot represents one gene set and is plotted according to its enrichment score (s) within the cell line dataset and the biopsy dataset. Significantly enriched (False Discovery Rate < 0.05) gene sets are shown in red.

Differential gene expression analysis revealed nearly twice as many DEGs between DFT1 and DFT2 cell lines compared to biopsies (8,612 DEGs and 4,569 DEGs, respectively). Cell lines show 4,200 up-regulated and 4,412 down-regulated genes in DFT2 compared to DFT1, while biopsies show 2,536 up-regulated and 2,033 down-regulated genes. Overall, more genes were detected in the cell lines than in the biopsies (17,468 and 15,931). Heatmaps of the top 20 up-and down-regulated genes, ranked by log fold-change (logFC) show cell lines and biopsies clustering by DFT (Fig. 1D). This grouping is less striking for the DFT2 biopsies, where 13 of the top 20 upregulated genes are solely overexpressed in one of the two samples. The most upregulated gene in DFT2 cell lines was *PDGFRA*, a receptor for platelet derived growth factors, and the most downregulated one was *NTS*, which may be involved in the regulation of fat metabolism [54]. In DFT2 biopsies, these genes are *SMTNL1* and *UBE2M*, respectively (see Table S2 for a full list of gene symbols, names, and brief descriptions). However, these genes are solely differentially expressed in one of the two biopsy samples.

Of the genes differentially expressed in both cell lines and biopsies, it was found that RFX4 and HOXD13, both of which encode transcription factors, and TNR, whose product is a neural extracellular matrix protein, were upregulated in both DFT2 cell lines and biopsies. Three genes, *CHRNA3*, *CLDN19* and *SSX2IP* were present within the top 20 downregulated genes in DFT2, both in cell lines and biopsies. CHRNA3 plays a role in neurotransmission, CLDN19 in tight junctions between cells and SSX2IP in eliminating ROS.

To understand pathway-level gene expression patterns, a multi-contrast enrichment analysis revealed 178 significantly enriched gene sets, i.e., sets with coordinated up or coordinated down regulation of genes (FDR < 0.05; Fig. 1E; Fig. S1). Most sets (153) showed concordant enrichment between cell line and biopsy data (top-right and bottom-left quadrants in Fig. 1E).

From these, 145 were upregulated in both DFT2 cell lines and biopsies and 8 were downregulated. Conversely, 25 significantly enriched gene sets showed discordant enrichment between cell lines and biopsies. Overall, although DFT cell lines diverge more in terms of gene expression than DFT biopsies, a vast majority of significantly enriched pathways are shared between both datasets.

Gene sets expected (or known) to be differentially expressed between DFT2 and DFT1 were concordantly upregulated in our analysis (Fig. S1): interferon gamma signalling, antigen presentation: folding, assembly and peptide loading of Major Histocompatibility Complex (MHC) class I, signalling by PDGF, nervous system development, axon guidance and somitogenesis [25, 26, 52, 55].

### Cholesterol biosynthesis and nucleotide metabolism show concordant enrichment in DFT2

Among the 178 enriched gene sets, a total of 14 pathways falls under the “metabolism” Reactome hierarchy (Fig. 2A). Two pathways showed concordant enrichment in DFT2 cell lines and biopsies: cholesterol biosynthesis (s_biopsies_ = 0.16 and s_cell lines_ = 0.47; Fig. 2B) and metabolism of nucleotides (s_biopsies_ = 0.07 and s_cell lines_ = 0.23; Fig. 2C). The cholesterol metabolism genes *SQLE*, *LSS* and *MVD* were upregulated in DFT2, in both datasets (Fig. 2B). Similarly, the *AK5*, *GMPR*, *TYMP*, *ENTPD4*, *NUDT18*, *IMPDH1*, *TXNRD1* and *GSR* genes, linked to the metabolism of nucleotides, were upregulated in DFT2 biopsies and cell lines. Within the same gene set, DFT2 cell lines and biopsies saw downregulated *AK9*.

**Figure 2.**
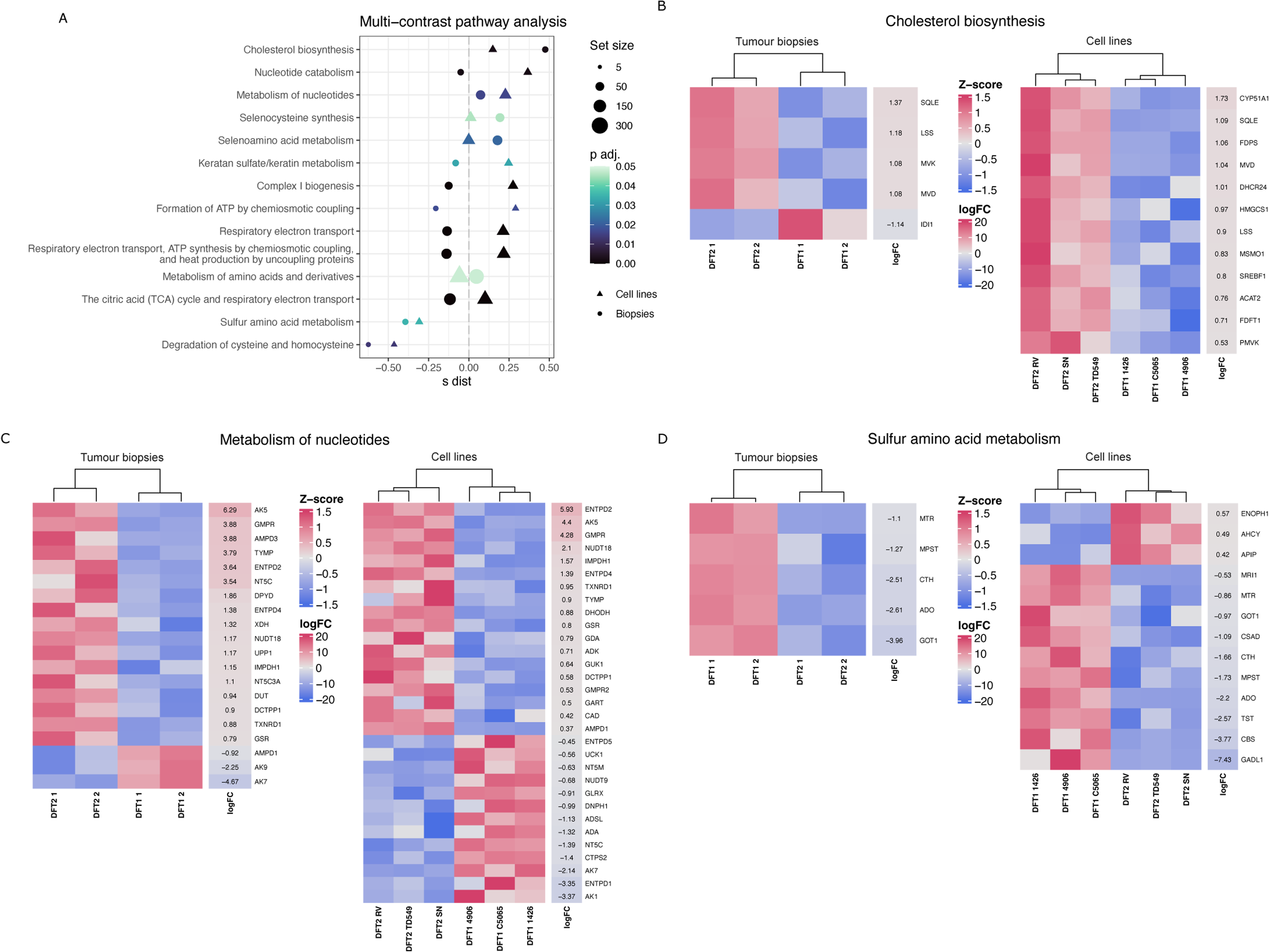
Cholesterol biosynthesis and nucleotide metabolism show concordant enrichment in Devil Facial Tumour 2 (DFT2). **A** Multi-contrast pathway analysis performed with mitch on Reactome pathways, filtered for metabolism-related gene sets with an adjusted pvalue < 0.05. Each circle or triangle represents one gene set and is plotted according to its enrichment score (s-dist). Triangle or circle size is scaled according to the number of genes in a pathway (set size). Heatmaps of Differentially Expressed genes (DEGs) for **B** cholesterol biosynthesis genes, **C** nucleotide metabolism genes and **D** sulfur amino acid metabolism genes. Only significant DEGs are represented (False Discovery Rate < 0.05).

Two metabolism related gene sets were concordantly enriched in DFT1 cell lines and biopsies: sulfur amino acid metabolism (s_biopsies_ = -0.31 and s_cell lines_ = -0.38) and degradation of cysteine and homocysteine (a subset of the sulfur amino acid metabolism gene set; s_biopsies_ = -0.46 and s_cell lines_ = -0.60; Fig. 2A). Sulfur amino acid genes, including *MTR*, *MPST*, *CTH*, and *ADO*, were downregulated in DFT2 cell lines and biopsies (Fig. 2D).

### DFT2 cells have higher glycolysis rates than DFT1 cells

Cholesterol biosynthesis was observed to be enriched in DFT2 cells and biopsies. It is known from previous research that cholesterol homeostasis is important for DFT1 cell proliferation [16] and can stimulate a metabolic shift towards glycolysis. This prompted us to initially quantify the relative contribution of glycolysis in DFT1 and DFT2 cell proliferation. Lactate, a by-product of glycolysis, was firstly measured in the cell culture medium (Fig. 3A). DFT2 cells tended to produce more lactate than DFT1 cells, with two DFT2 cell lines (SN, TD549) showing significantly increased extracellular lactate production over time. SN produces 30.51 nmol of lactate, 95% CI [26.76, 34.59], whilst the highest lactate production by a DFT1 cell line was 2.7 nmol (p < 0.01; 95% CI [0.13, 11.85]). Remarkably, the DFT2 TD549 cell lines produced over a hundred times more lactate (174.83 nmol, 95% CI [139.51, 215.68]) compared to other cell lines. This higher lactate production by DFT2 cells indicates higher rates of glycolysis compared to the DFT1 cell lines.

**Figure 3.**
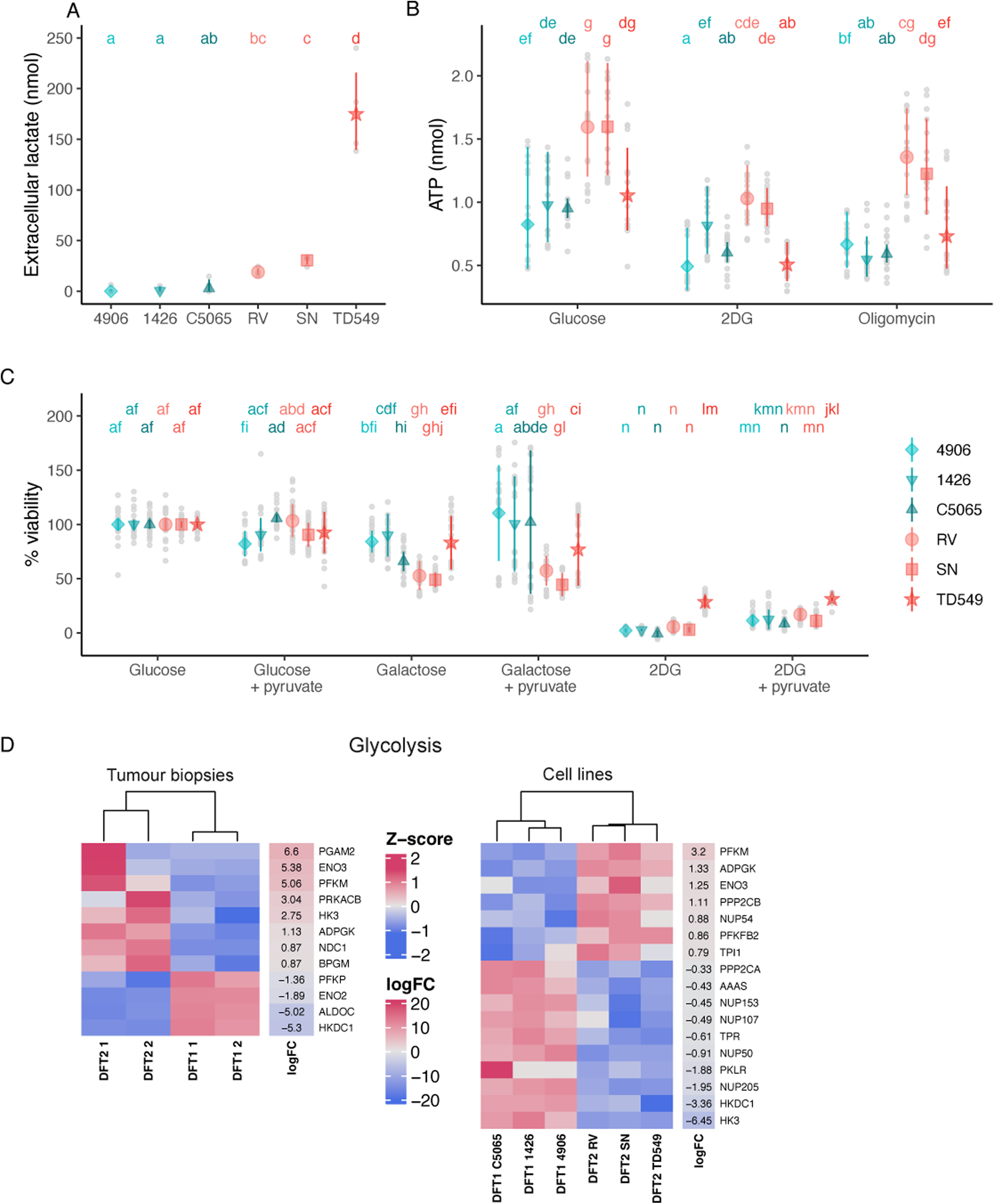
Devil Facial Tumour 2 (DFT2) cells show higher glycolysis rates than DFT1 cells. **A** Lactate quantity measured in cell culture supernatant, best fit model: lactate ∼ cell line + day. Two independent experiments were performed, each comprised duplicates for each cell line. **B** Total (glucose), oxidative (2-deoxy-D-glucose, 2DG) and glycolytic (oligomycin) adenosine triphosphate (ATP) production. Best fit model: ATP ∼ treatment * cell line + (1 | experiment). Three independent experiments were performed, each comprised of 6 technical replicates per cell line. **C** Percent viability of Devil Facial Tumour 1 (DFT1, blue) and DFT2 (red) cell lines grown in treatments targeting glucose metabolism relative to glucose control medium. Best fit model: % viability ∼ treatment * cell line + (1 | experiment). Three independent experiments were performed; each comprised 6 technical replicates per cell line. **A**, **B** and **C** Mean and 95 % confidence intervals shown in colour and data points in grey. Compact letter display represents post hoc contrasts for which p < 0.05. Following Tukey’s post hoc test (with Bonferroni correction) each group was assigned a letter. Groups that share a letter are not significantly different from each other. Groups with different letters are significantly different at a threshold of p < 0.05. **D** Heatmaps of DEGs for glycolysis genes. Only significant DEGs are represented (False Discovery Rate < 0.05).

We then tested how this difference in glycolysis rates would impact energy production in the form of ATP (Fig. 3B). To compare the amount of energy produced by glycolysis versus OXPHOS, we reported total, oxidative and glycolytic ATP production. Two DFT2 cell lines overall produced 1.5 times more ATP than DFT1 cells when cultured in media with glucose only; RV produced 1.59 nmol of ATP (95% CI [1.27, 2.01]) and SN cells 1.6 nmol (95% CI [1.28, 2]), compared to the highest ATP production in DFT1 cells of only 0.98 nmol by 1426 (p < 0.001, 95% CI [0.73, 1.31]) (Fig. 3B).

Exposure of cells to 2-deoxyglucose (2DG, which is converted to 2-deoxyglucose-6-phosphate (2DG6P) and accumulates in the cell, ultimately inhibiting glycolysis [56]), significantly decreased oxidative ATP production between 0.33 to 0.65 nmol in all cell lines (except one DFT1 cell line, 1426) compared to media with glucose only (p < 0.001) (Fig. 3B). Inhibiting OXPHOS with oligomycin (glycolytic ATP production only) significantly decreased ATP production in two DFT1 cell lines (1426 by 0.43 nmol and C5065 by 0.36 nmol; p < 0.001), and one DFT2 cell line (TD549 by 0.32) compared to glucose only treatment (Fig. 3B), In summary, DFT2 cell lines have higher rates of glycolysis and ATP production that DFT1 cell lines and rely less on OXPHOS for ATP generation than DFT1 cells. Furthermore, DFT1 cells are more sensitive to inhibition of either glycolysis or OXPHOS for ATP generation than DFT2 cells. Interestingly, the DFT2 TD549 cell line appears to rely more on oxidative ATP generation than the other DFT2 cell lines (i.e. more sensitive to oligomycin). In conjunction with its high rate of lactate generation and lower ATP levels in media with glucose only, this finding suggests that ATP generation is less efficient overall in TD549 cells.

To further investigate the importance of metabolic selection in DFT1 and DFT2, we measured cell viability in the presence of various substrates (Fig 3C). Both DFT1 and DFT2 showed a significant decrease in cell viability by at least 71.6 % (p < 0.001) when glycolysis was inhibited with 2DG, compared to growth in glucose alone (Fig 3C). To test whether cells could switch to OXPHOS to obtain the ATP necessary for their survival, glucose was replaced by galactose. Galactose can enter glycolysis after being converted to glucose-6-phosphate through the Leloir pathway. However, this is less efficient and results in no net ATP gain [57]. Galactose significantly decreased viability in one DFT1 cell line (C5065 by 34.09 %; p < 0.001) and in two DFT2 cell lines (RV by 47.28 % and SN by 51.09 %; p < 0.001).

Lastly, pyruvate was added to the 2DG and galactose treatments to assess whether this could restore cell viability by driving OXPHOS [58]. Pyruvate did not rescue viability in any cell line when added in conjunction with 2DG. In contrast, when added in conjunction with galactose, pyruvate significantly increased the survival of two DFT1 cell lines (4906 by 26.33 % and C5065 by 36.22 %; p < 0.001) compared to galactose only treatment (Fig 3C).

In summary, both DFT1 and DFT2 cell lines rely on glycolysis for survival. However, DFT1 appears more metabolically flexible and can switch to oxidative metabolism (OXPHOS) when required, whereas DFT2 cannot. Interestingly, the TD549 DFT2 cell line showed significantly higher survival compared to the other two DFT2 cell lines in most treatments (by at least 30.47 % in galactose, p < 0.001; 19.58 % in galactose and pyruvate, p < 0.01; and 22.8 % in 2DG, p < 0.001), suggesting these cells are more metabolically flexible than the other DFT2 cell lines examined.

Linking the functional and transcriptomic data, we identified genes that could be responsible for DFT2’s higher glycolytic rates. Although the glycolysis gene set was not enriched in DFT2 (p > 0.05), three glycolysis genes were upregulated in DFT2: *ENO3*, *PFKM* and *ADPGK* (Fig. 3D). One gene, *HKDC1*, was downregulated in DFT2 in both datasets, whereas *HK3* was upregulation in DFT2 biopsies but was downregulated in DFT2 cell lines. This suggests that *ENO3*, *PFKM* and *ADPGK* are primarily driving the increase in glycolytic rates observed in DFT2 compared to DFT1.

### DFT2 cells produce more ROS than DFT1 cells

We next investigated the effect of increased glycolysis and ATP production on ROS generation (Fig. 4A), as higher levels of glycolysis have been associated with elevated ROS production and oxidative stress [59]. Overall, ROS production was significantly higher in DFT2 cell lines, with RV producing the highest amount of ROS (70624 RFU, 95% CI [23282, 31874], p < 0.001), followed by SN (39949 RFU, 95% CI [32283, 47615] compared to DFT1 lines 1426 (22679, 95% CI [18272, 27087], p < 0.01) and C5065 (12439 RFU, 95% CI [9479, 15400], p < 0.01). To identify the type of ROS being produced, we quantified mitochondrial superoxide ROS production (Fig. 4B), however, no significant difference in the production of mitochondrial ROS was observed between any cell line (p > 0.05). DFT2 thus produces more ROS than DFT1, with this increase being cytoplasmic (and not mitochondrial) in origin.

**Figure 4.**
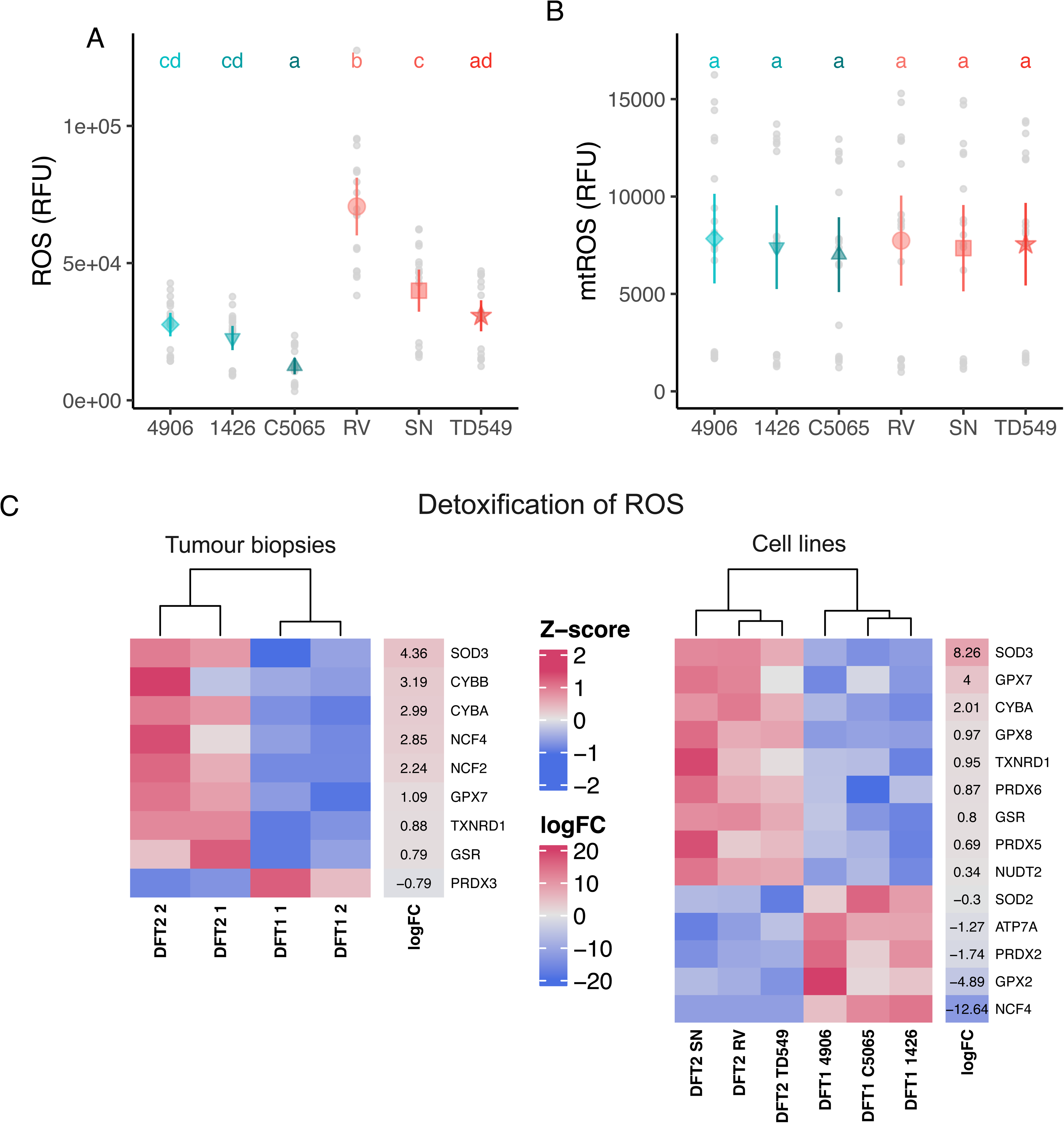
Devil Facial Tumour 2 (DFT2) cells produce more reactive oxygen species (ROS) than DFT1 cells. **A** Overall ROS and **B** mitochondrial ROS production in relative fluorescence units (RFU) of DFT1 and DFT2 cell lines after 1 hour. Best fit model: ROS ∼ cell line. Three independent experiments were performed; each comprised 6 technical replicates per cell line. Mean and 95 % confidence intervals shown in colour and data points in grey. Compact letter display represents post hoc contrasts for which p < 0.05. Following Tukey’s post hoc test (with Bonferroni correction) each group was assigned a letter. Groups that share a letter are not significantly different from each other. Groups with different letters are significantly different at a threshold of p < 0.05. **C** Heatmaps of differentially expressed genes (DEGs) for ROS detoxification genes. Only significant DEGs are represented (False Discovery Rate < 0.05).

In line with the observed elevated ROS production, both DFT2 cell lines and biopsies show enrichment of ROS detoxification genes (Fig. 4C; Fig. S1). *SOD3*, *CYBA*, *GPX7*, *TXNRD1* and *GSR* are overexpressed in both DFT2 datasets. Interestingly, *NCF4* is upregulated in DFT2 biopsies but downregulated in DFT2 cell lines. This suggests that increased cytoplasmic ROS production by DFT2 is somewhat mitigated by the upregulation of ROS detoxification genes to avoid ROS-induced cell death.

### DFT2 relies more on lipid metabolism than DFT1

Lipid metabolism is essential in rapidly dividing cancer cells [60], which led us to test the role of fatty acid metabolism in DFT1 and DFT2 proliferation. We used etomoxir, an irreversible carnitine palmitoyl-transferase 1A inhibitor (CPT1A), to inhibit fatty acid oxidation [61] (Fig. 5A). Etomoxir acts by causing lipid accumulation and decreased ATP and NADPH production, ultimately stopping cell proliferation [62]. Etomoxir significantly decreased the viability of two DFT2 cell lines (RV by 35.34 % and SN by 38.17 %; p < 0.001), but not in TD549 (by 1.58 %; p > 0.05) compared to glucose only controls. Conversely, DFT1 cell lines had no change in survival when cultured in etomoxir compared to glucose only controls (Fig. 4A).

**Figure 5.**
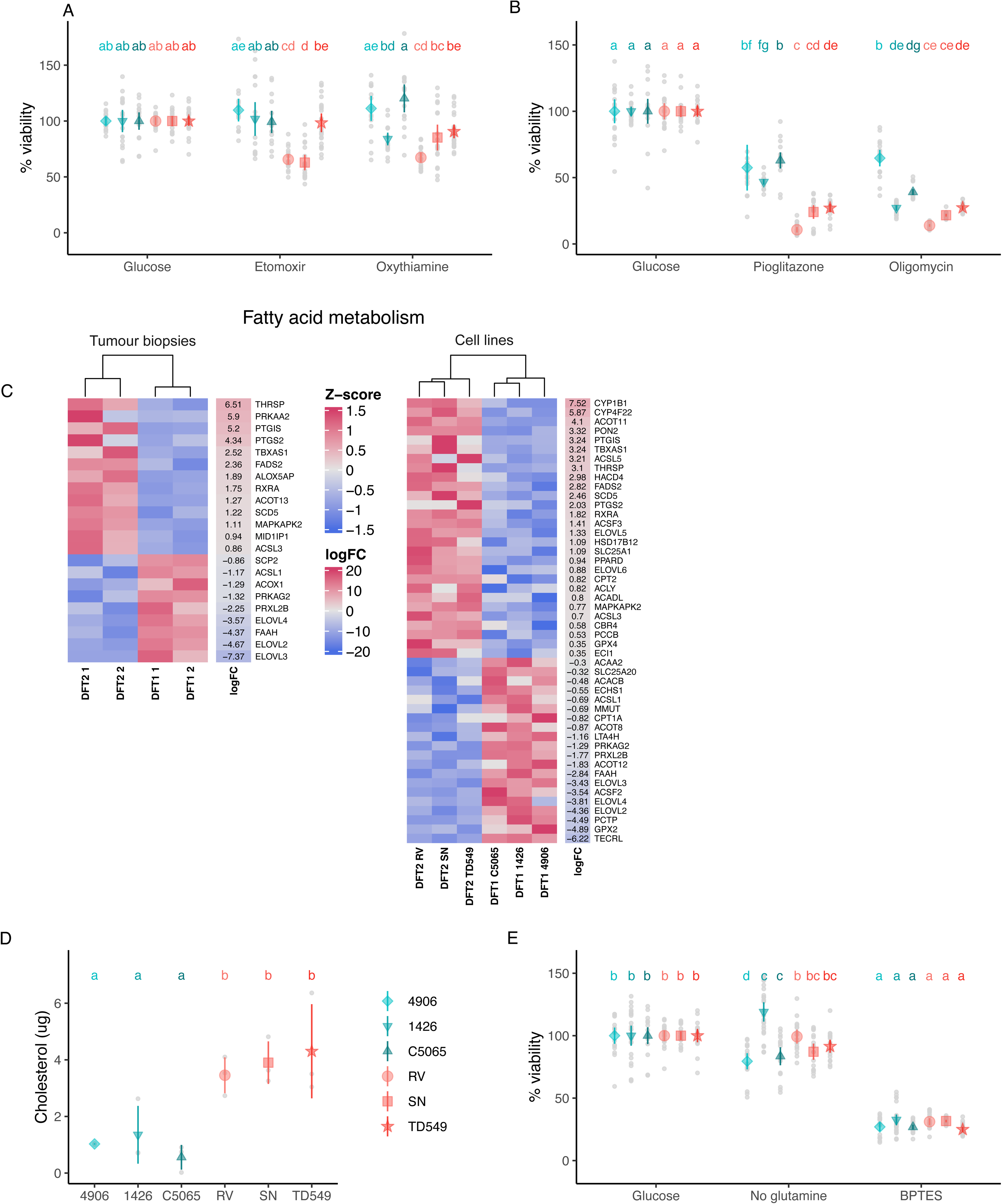
Devil Facial Tumour 2 (DFT2) relies more on lipid metabolism than DFT1, and other pathways of interest. Percent viability of DFT1 (blue) and DFT2 (red) cell lines grown in treatments targeting **A** fatty acid oxidation and the pentose phosphate pathway (PPP) and **B** fatty acid oxidation and oxidative phosphorylation (OXPHOS). **C** Heatmaps of differentially expressed genes (DEGs) for fatty acid metabolism. Only significant DEGs are represented (False Discovery Rate < 0.05). **D** Cholesterol content of lipids extracted from 10^6^ cells, best fit model: cholesterol ∼ DFT. Three samples per cell line, each from independent cell subcultures, were used for this experiment. **E** Percent viability of DFT1 and DFT2 cell lines grown in treatments targeting glutamine metabolism. **A**, **B** and **E** Three independent experiments were performed; each comprised 6 technical replicates per cell line. Best fit models: % viability ∼ treatment * cell line. **A**, **B**, **D** and **E** Mean and 95 % confidence intervals shown in colour and data points in grey. Compact letter display represents post hoc contrasts for which p < 0.05. Following Tukey’s post hoc test (with Bonferroni correction) each group was assigned a letter. Groups that share a letter are not significantly different from each other. Groups with different letters are significantly different at a threshold of p < 0.05.

We also investigated the importance of the pentose phosphate pathway (PPP) for providing NADPH (as well as nucleotides) for DFT cell viability [63] (Fig. 5A). For these experiments we used oxythiamine to prevent thiamine from transitioning to its active form, ultimately inhibiting transketolase, an essential enzyme in the PPP [64]. Oxythiamine had no effect on DFT1 survival, but showed a trend of reduced viability in DFT2 cells, with viability significantly decreased in the RV DFT2 cell line (by 32.66 %; p < 0.001).

To investigate if DFTs also rely on fatty acid anabolism, cells were further treated with pioglitazone, an ACSL4 inhibitor blocking fatty acid synthesis [61] (Fig. 5B). When treated with pioglitazone, all DFT cell lines showed decreased viability compared to untreated glucose only cells (p < 0.001). Pioglitazone had a greater antiproliferative effect on DFT2 cell lines (by 89.29 %), compared to DFT1 (by 35.33 %; p < 0.001). We hence conclude that DFT2 cells rely more on both fatty acid oxidation and fatty acid synthesis than DFT1 cells.

We also examined if DFT1 and DFT2 cells rely on oxidative ATP generation for their viability (Fig. 5B). DFT cells were treated with the ATP synthase inhibitor oligomycin, which significantly decreased cell viability for all six cell lines compared to untreated (glucose only) controls (Fig 5B; by 42.5% on average, p < 0.001). This demonstrates that oxidative phosphorylation, and the ATP generated by this pathway, remains important for DFT cell survival.

Linking the fatty acid metabolism results with the transcriptomics data, we found that the following fatty acid metabolism genes were concordantly upregulated in DFT2 cell lines and biopsies compared to DFT1: *THRSP*, *PTGIS*, *PTGS2*, *TBXAS1*, *FADS2*, *SCD5*, *MAPKAPK2* and *ACSL3* (Fig. 5C). Conversely, *ACSL1*, *PRKAG2*, *PRXL2B*, *ELOVL4*, *FAAH*, *ELOVL2* and *ELOVL3* were downregulated in DFT2 compared to DFT1. It is important to note however that the total fatty acid metabolism gene set is not significantly enriched in DFT2 (p > 0.05).

Based on our previous observation that cholesterol biosynthesis genes are enriched in DFT2, we measured cholesterol content in DFT1 and DFT2 cells (Fig. 5D). DFT2 cells contained three to four times more cholesterol than DFT1 cells (p < 0.05), consistent with DFT2 upregulating genes linked to cholesterol biosynthesis (Fig. 2A & B).

### Both DFT1 and DFT2 rely on glutamate production and oxidative metabolism for survival

DFT1 and DFT2’s reliance on glutamine, an essential substrate for cancer cell metabolism [65], was tested by incubating cells in glutamine depleted medium (Fig. 5E). This significantly decreased the viability of only one DFT1 cell line (4906 by 20.5 %; p < 0.01) and significantly increased the viability of another DFT1 cell line (1426 by 19.03 %; p < 0.001) compared to control medium with glucose (and glutamine). DFT2 cell lines were not impacted by the absence of glutamine in their culture medium. The DFT1 and DFT2 cells were then grown with BPTES, an inhibitor preventing glutaminase-1 from converting glutamine to glutamate, and this treatment significantly decreased the viability of all DFT1 and DFT2 cell lines compared to control glucose (with glutamine) treatment (by 72.99 % on average; p < 0.001). This result suggests that DFT1 and DFT2 can generate endogenous glutamine, but both rely on the conversion of this glutamine to glutamate for survival.

## DISCUSSION

This work demonstrates that, *in vitro*, similar metabolic pathways drive DFT1 and DFT2 proliferation, with glycolysis, OXPHOS, glutamine metabolism and fatty acid synthesis being essential for their survival. However, the two transmissible tumours differ slightly, but significantly, in their metabolic usage. DFT2 has a higher rate of glycolysis relative to DFT1, which is accompanied by higher lactate generation and ATP production. This is consistent with DFT2’s hypothesised cell-type of origin (i.e., a repair Schwann cell, whilst DFT1 originated from a myelinating Schwann cell [26, 66]). During nerve injury, repair Schwann cells increase glycolysis in comparison to myelinating Schwann cells to support axons [27]. Our transcriptomic analysis revealed that DFT2 could achieve this by upregulating *ADPGK*, *PFKM*, and *ENO3*. ADPGK catalyses phosphorylation of glucose to glucose-6-phosphate using ADP as a phosphate donor. This initial step of glycolysis is typically catalysed by ATP-dependent hexokinases. Therefore, it is believed that ADPGK could assist in conserving ATP during hypoxia, thereby reducing the ATP consumption necessary for glucose phosphorylation [67]. Next, PFKM catalyses the phosphorylation of D-fructose 6-phosphate to fructose-1,6-bisphosphate using ATP, which is the first committed step of glycolysis. Finally, ENO3 is responsible for the conversion of 2-phosphoglycerate to phosphoenolpyruvate in the glycolytic pathway. Because fructose-1,6-bisphosphate production has previously been identified as one of the key flux-controlling steps of glycolysis [68], we expect *ADPGK* and *PFKM* upregulation to play a major role in DFT2’s higher rate of glycolysis in comparison to DFT1. Interestingly, we also observed upregulation of hexokinase 3 (*HK3*) in DFT2 biopsies, which further suggests this could be a mechanism underlying higher rates of glycolysis.

This is the first study directly comparing DFT1 and DFT2 metabolism using tumour cell lines and biopsies. At the time of this work, DFT2 transcriptomes were scarce with only a single study conducting full RNA sequencing of one DFT1 and DFT2 cell line and two DFT1 and DFT2 biopsies [26]. By conducting full transcriptome analyses of two additional DFT1 and DFT2 cell lines, this study provides valuable data for future *in vitro* work on DFTD. Overall, more genes were differentially expressed in the cell lines than in biopsies, likely due to differences in sequencing depth between the transcriptomes. In the future, re-sequencing both cell lines and biopsies on the same platform would offer a more robust approach, as this difference could have led to an underestimation of the number of differentially expressed genes in the biopsy data. Nevertheless, by integrating *in vitro* metabolic assays with the analysis of cell line and biopsy transcriptomes, we minimize the risk of identifying metabolic differences associated with adaptation to cell culture conditions. Consequently, we can confidently identify metabolic pathways crucial for these transmissible tumours.

Previous research [16] had demonstrated the importance of glycolysis for DFT1 proliferation. We confirm their findings by showing the reliance of both DFT1 and DFT2 on glycolysis for survival. Ikonomopoulou and colleagues [16] further linked cholesterol homeostasis to this metabolic shift towards glycolysis in DFT1. We observe a similar pattern in DFT2. Cholesterol biosynthesis genes are enriched in DFT2 cell lines and biopsies, which is consistent with DFT2 containing threefold more cholesterol than DFT1 and its higher rate of glycolysis. Indeed, the acidic environment caused by elevated glycolysis and ATP hydrolysis has been shown to increase cholesterol biosynthesis in human malignancies, allowing cancer cells to sustain proliferation, migrate and invade tissues [69].

Another finding of our study is that DFT2 produces more cytoplasmic ROS than DFT1. Current research [59] shows that increased glucose conversion to pyruvate can elevate cellular ROS levels. We thus hypothesise that higher glycolysis rates driven by *ADPGK* and *PFKM* upregulation could be responsible for both increased ROS production and upregulation of ROS detoxification genes in DFT2. Furthermore, we observed concordant enrichment of genes belonging to the degradation of cysteine and homocysteine pathway in DFT1 cell lines and biopsies. Degradation of these amino acids through the transulphuration pathway leads to the generation of glutathione (GSH) and taurine, both sustaining redox chemistry [70]. This could explain lower ROS production by DFT1 compared to DFT2 cell lines, although this needs to be verified by quantifying GSH production.

Within the DFT2 cell lines tested in this study we mostly observed comparable effects to the inhibitors used, although TD549’s response tended to differ from the other two cell lines (RV and SN). As such, TD549 produced less ATP and ROS, more lactate, and had an overall better survival in the presence of inhibitors than the other DFT2 cell lines. We hypothesise that rearrangements on chromosome 2 distinguishing TD549 from RV and SN could explain such differences [71]. TD549 notably lacks one copy of the gene *FSHR* [71], the product of which is responsible for downstream PI3K-AKT signalling [72], a pathway implicated in altered carbohydrate metabolism in DFT1 [16]. In Figure 1A the multidimensional scaling (MDS) analysis reveals that the biological replicates of DFT1 tumours cluster together in a distinct pattern separate from the DFT2 tumours. Interestingly, the DFT2 biological replicates also cluster separately from each other. This difference in DFT2 tumour clustering could also be influenced by factors associated with the original tumour biopsy, including the stage of the cancer, its specific location and possible tumour cell sub-linages.

Our findings are somewhat limited due to the use of *in vitro* conditions that do not accurately reflect the genetic and vascular heterogeneity of an *in vivo* tumour [73]. In addition, one might not capture the full extent of the diversity of tumours circulating currently in Tasmania, a problem which could be addressed by deriving more recent cell lines from primary tumours.

Increased glycolysis rates in cancer cells play an important role in immune evasion by limiting the availability of glucose to tumour-infiltrating lymphocytes [8]. DFT1 is thought to evade the immune system by downregulating MHC molecules, which are still expressed by DFT2 [55, 74]. Furthermore, certain DFT2 lineages carry a Y chromosome (as DFT2 originated from a male devil) and, likely because of its immunogenicity, appear to be less successful at infecting female devils [22, 75]. Therefore, we hypothesise that increased lactate production in DFT2, because of increased glycolysis rates, might play a role in evading the devil’s immune system. Indeed, previous research reported that few tumour-infiltrating lymphocytes were observed within DFT2 biopsies [55]. That would explain the differential gene expression patterns between the two DFT2 biopsy samples (Fig.1D), as two B cell and plasma cell markers (MS4A1 and MZB1) were highly expressed in DFT2_2 biopsy, indicating immune infiltration. An active immune response in a tumour would consume a lot of energy, and thus potentially impact the expression of metabolism related genes for that sample. As our tumour purity calculations indicated the sample mostly consisting of tumour cells and as we focused on genes that were differentially expressed/pathways enriched in both the biopsies and in the cell lines, we believe the general conclusions remain the same.

ROS are a source of oxidative DNA damage and result in higher DNA mutation rates. We observed higher ROS generation in DFT2 cell lines compared to DFT1, and this concurs with the higher mutation rates observed in DFT2 tumours compared to DFT1 tumours [22]. High mutation rates in clonally reproducing organisms should lead to the accumulation of deleterious mutations, and eventually genomic decay [76, 77]. However, the three types of transmissible cancers known so far all share mutation rates exceeding those of human cancers, and it is proposed that such a high mutation load could carry indirect positive fitness outcomes [78].

DFT2’s higher rate of glycolysis explains its previously reported faster *in vitro* growth and could explain how it is able to outcompete DFT1 in co-culture assays [30]. Higher rates of cell division require increased deoxyribonucleotide triphosphate (dNTP) synthesis, which are not only rate-limiting for DNA replication and repair, but also for transcription, ribosome biogenesis and post-translational protein glycosylation [79]. Indeed, we found that reducing nucleotide supply by inhibiting the pentose phosphate pathway decreased DFT2 cell viability (Fig 5A). This corresponds with the enriched nucleotide metabolism and catabolism in DFT2 cells that drive faster cell division [30].

Considering these observations, we hypothesise that DFT2 could be a more aggressive cancer than DFT1. Meta-analyses of human cancer patients have associated elevated aerobic glycolysis with poorer prognosis [80, 81]. For instance, elevated expression of glycolysis-related genes was associated with aggressive phenotypes such as poor tumour differentiation, deeper invasion, lymph node metastasis and vascular invasion in high-grade brain tumours [81]. A second meta-analysis [80] demonstrated ultimately worse overall survival for hepatocellular carcinoma patients with elevated expression of glycolysis markers. Based on these human studies, it can be predicted that DFT2 tumours could have a higher potential to invade tissues and metastasise.

The fact that DFT1 and DFT2 appear to share some common pathways for their growth and survival is promising for the development of future treatments against DFTD. Atorvastatin, which has proven to be effective in reducing DFT1 tumour growth [16], is likely to also inhibit DFT2 tumour growth, underscoring its potential to assist in ongoing management plans for the conservation of Tasmanian devils. However, DFT2’s increased glycolytic rate and potential metabolic plasticity could represent an important threat to Tasmanian devils if these metabolic features translate *in vivo*. If so, DFT2 is likely to have a faster growth rate, enhanced immune evasion and higher adaptive potential than DFT1. It is important to note that DFT2 has been progressing on a different epidemiological landscape than DFT1. DFT2 arose in a smaller, more disperse population that had already been exposed to DFT1 for almost 30 years . However, DFT2’s evolutionary trajectory remains unknown. Our results, taken together with DFT2’s expansion outside of the peninsula where it originated, highlight the necessity to further research the differences and similarities of these unique transmissible cancers and their endangered host.

Finally, to survive across hosts and environments, we show that Tasmanian devil facial tumour cells undergo extreme metabolic flexibility, which mirrors what human cancers do during metastasis and therapy resistance. Like human tumours, transmissible cancers rely heavily on glycolysis—even when oxygen is available, and such Warburg effect appears to be amplified in these clonal cells. They exhibit metabolic plasticity as they adapt their energy sources depending on the host’s tissue environment, showcasing how tumours can switch between glycolysis, oxidative phosphorylation, and lipid metabolism. Transmissible cancers offer real-world examples on how cancer cells survive in hostile or changing environments. The pathways identified in our study reveal the metabolic vulnerabilities of these, and potentially human cancers, offering druggable targets and treatment options. As DFTD cells use similar metabolic strategies as human cancers, they can serve as natural models to test metabolic inhibitors (e.g., drugs targeting glycolysis, glutamine metabolism, or lipid synthesis) and may be used to help identify biomarkers for aggressive or treatment-resistant human cancers. While transmissible cancers are rare, they can show how cancer cells can thrive, evolve, and adapt metabolically, and such provide a window into the metabolic character of cancer.

## MATERIALS AND METHODS

### EXPERIMENTAL MODEL

#### Cell Lines and Cell Culture

Devil Facial Tumour 1 (DFT1) cell lines 4906, 1426 and C5065 and DFT2 cell lines RV, SN and TD549 were grown *in vitro* as previously described [21, 30, 31]. DFT cell lines were derived from different animals and act as biological replicates. Cells were cultured in RPMI-1640 media (Gibco, 21870092; 2 g/L of glucose), supplemented with 1 % GlutaMAX (Gibco, 35050061), 10 % FBS (Gibco, 10099141) and 1 % 10,000 U/mL penicillin/streptomycin (Gibco, 15070063) at 35 °C and 5 % CO_2_. Upon reaching 80-90 % confluency, cells were washed with DPBS (Gibco, 14190250) detached using TrypLE Express (Gibco, 12605010), centrifuged at 500 x g for 5 minutes and passaged 1:3. Cells used in the present work were maintained below passage 30. All cell lines were obtained from Dr Hannah V. Siddle. DFT1 cell lines were tested for mycoplasma as described in [29]. DFT2 cells were tested for mycoplasma using the MycoAlert mycoplasma testing kit (Lonza, LT07-418) when they entered the laboratory and were passaged in a mycoplasma-free tissue culture facility after testing [30]. Additional information on the cell lines used in the present study can be found in Table 1.

**Table 1.**
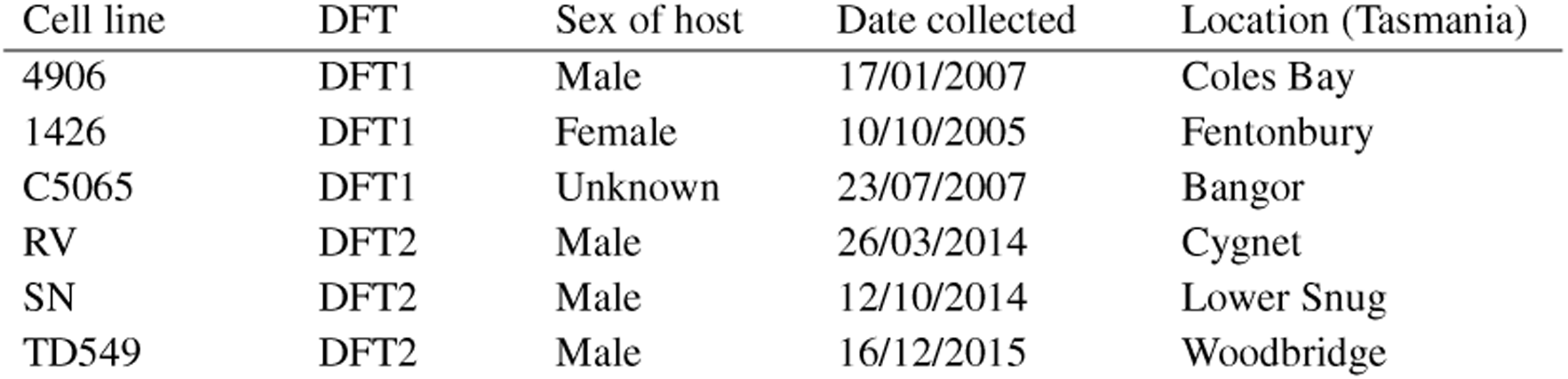
Cell lines used in the present study.

### METHOD DETAILS

#### Sample Processing and RNA Sequencing

DFT cell lines were grown in 6-well plates until 80-90 % confluent before harvesting. RNA was extracted from all samples using the Zymo Direct-zol™ RNA kit (Zymo Research, R2051) as per manufacturer instructions. Quantification and qualification of extracted RNA was performed on a Qubit RNA High Sensitivity (Invitrogen) and on a Tapestation 4200-RNA Screentape (Agilent). All samples had RNA integrity scores (RIN) scores greater than 9.5.

#### Library Preparation and Sequencing

Library preparation was performed using the NEBNext® Ultra™ II Directional RNA Library Prep Kit for Illumina (New England Biolabs, E7760) with 400 ng of each sample, 5X Adapter dilution and a 10 cycle PCR. Technical triplicates from all samples were sequenced at the Charles Rivers Laboratories on a NovaSeq6000 (Illumina) using 300 cycles and 150 paired end reads.

#### Previously published data

RNA sequences from two DFT1 and two DFT2 primary tumours published in Patchett et al. [26] were re-analysed for comparison will cell line data. These tumour data consist of two biological replicates per DFT, sequenced in duplicate (i.e., technical replicates) using 100 base-pair single end read mRNA sequencing on the Hiseq-2500 platform (Illumina). FASTQ files were downloaded from the European Nucleotide Archive (project PRJEB28680).

#### Metabolic assays

Cell viability after metabolic inhibition was measured by MTT (3-(4, 5-dimethylthiazolyl-2)-2, 5-diphenyltetrazolium bromide) assay (Cell Proliferation Kit I, Roche, 11465007001) following manufacturer’s instructions. 10,000 cells/well were seeded in 96-well and incubated for 24 h. Cells were then treated with fresh culture medium (glucose controls) or culture medium with metabolic inhibitor (treatments; see Table 2). Concentration of inhibitors was determined based on common concentrations used in the literature [32–37]. Cells were incubated for an additional 72 h before MTT reduction was measured at 550 nm in a FLUOstar Omega plate-reader (BMG LABTECH) and Omega software (version 3.00 R2). Controls were used to define 100 % viability with three independent experiments performed, each comprised of 6 technical replicates per cell line.

**Table 2.**
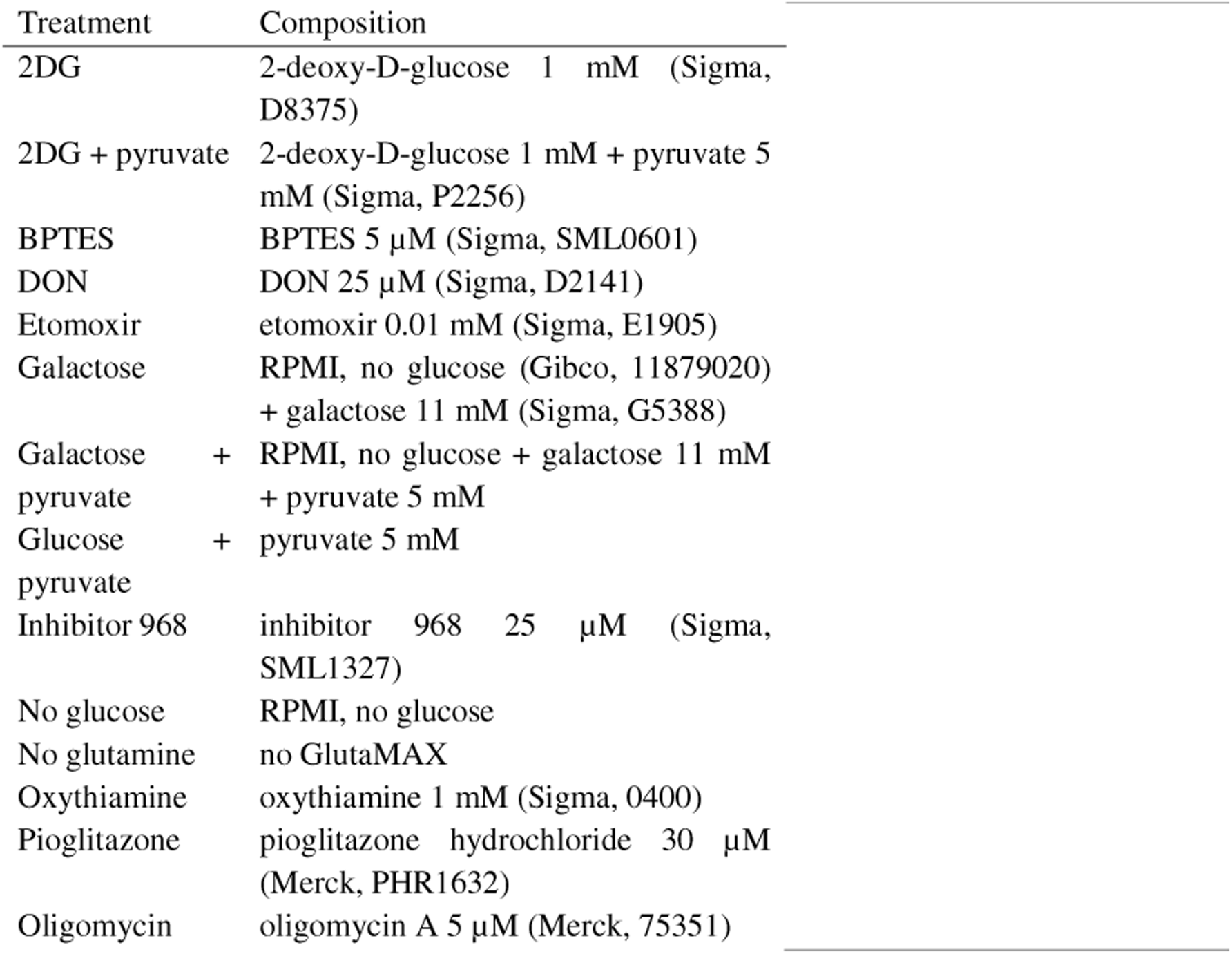
Treatment conditions and compositions in the metabolic inhibition experiments.

#### Measurement of ATP production

Cells were treated with fresh culture medium (control), culture medium with 1 mM 2DG (Sigma, D8375; to measure oxidative ATP production) or 5 μM oligomycin (Sigma, 75351; to measure glycolytic ATP production) as previously described [38] and incubated for 3 h. ATP was quantified using the ATPlite 1step kit (PerkinElmer, 6016731) according to manufacturer’s instructions. Luminescence was measured in a FLUOstar Omega plate-reader (BMG Labtech). Three independent experiments were performed, each comprised of 6 replicates per cell line.

#### Measurement of ROS production

Overall ROS production was measured using H_2_DCFDA (Sigma, D6883). Cells were incubated with 10 μM H_2_DCFDA for 30 min in the dark at 35°C, washed with DPBS, then incubated for 20 min in 100 μL of DPBS. Fluorescence at 480 nm excitation and 520 nm emission was measured using a FLUOstar Omega plate-reader for 1 h. Three independent experiments were performed, each comprised of 6 replicates per cell line.

Mitochondrial ROS production was quantified using MitoSOX™ Green (Thermo Fisher Scientific, M36006). Cell were seeded on black clear bottom 96-well plates (Interpath, 655090) and left to adhere for 48 h. Cells were then incubated with 1 µM MitoSOX™ Green in DPBS and incubated at 35°C and 5% CO_2_ for 30 min. Cells were washed and plates centrifuged at 400 x *g* for five min, followed by incubation for one h in the dark. Fluorescence intensity was then measured at 485 nm excitation/520 nm emission with a FLUOstar Omega plate-reader. Three independent experiments were performed with six replicates per cell line.

#### Measurement of extracellular lactate

Cells were plated and incubated as described above. Cell culture supernatants were harvested after 2, 6 and 8 days, deproteinised using 10 kDa MWCO spin filters (Merck, UFC5010), and stored at -80 °C until further analysis. After each collection, cell number per well was measured by MTT. Cell culture supernatants were thawed, and lactate was measured using a Lactate Assay Kit (Sigma, MAK064) according to the manufacturer’s protocol. Absorbance was measured at 570 nm on a FLUOstar Omega plate-reader. Wells containing only media were used to blank the readings. Two independent experiments were performed, each comprised of duplicates for each cell line. Lactate content was standardised by cell number and calculated for 10^6^ cells.

#### Measurement of cholesterol content

One million cells were pelleted and stored at -80 °C until further processing. Samples were thawed and were resuspended in 200 μL of a 7:11:0.1 chloroform:isopropanol:IGEPAL mix (Merck, 1.02445; Chem Supply, AH323-4; Merck, CA-630) to extract lipids. Samples were centrifuged for 10 min at 13,000 x g to remove insoluble material. The organic phase was transferred to a new tube and left to air dry at 50 °C for 1 h. Then samples were dehydrated under a vacuum for 30 min at 50 °C using a SpeedVac (Savant). Dried lipids were resuspended in 200 μL of Cholesterol Assay Buffer from the Cholesterol Quantitation Kit (Sigma, MAK043). Manufacturer’s directions were followed to measure cholesterol content. Absorbance was recorded at 570 nm on a FLUOstar Omega plate-reader. Three samples per cell line, each from independent cell subcultures, were used for this experiment.

Table 3 contains information on all key resources used in the study.

**Table 3.**
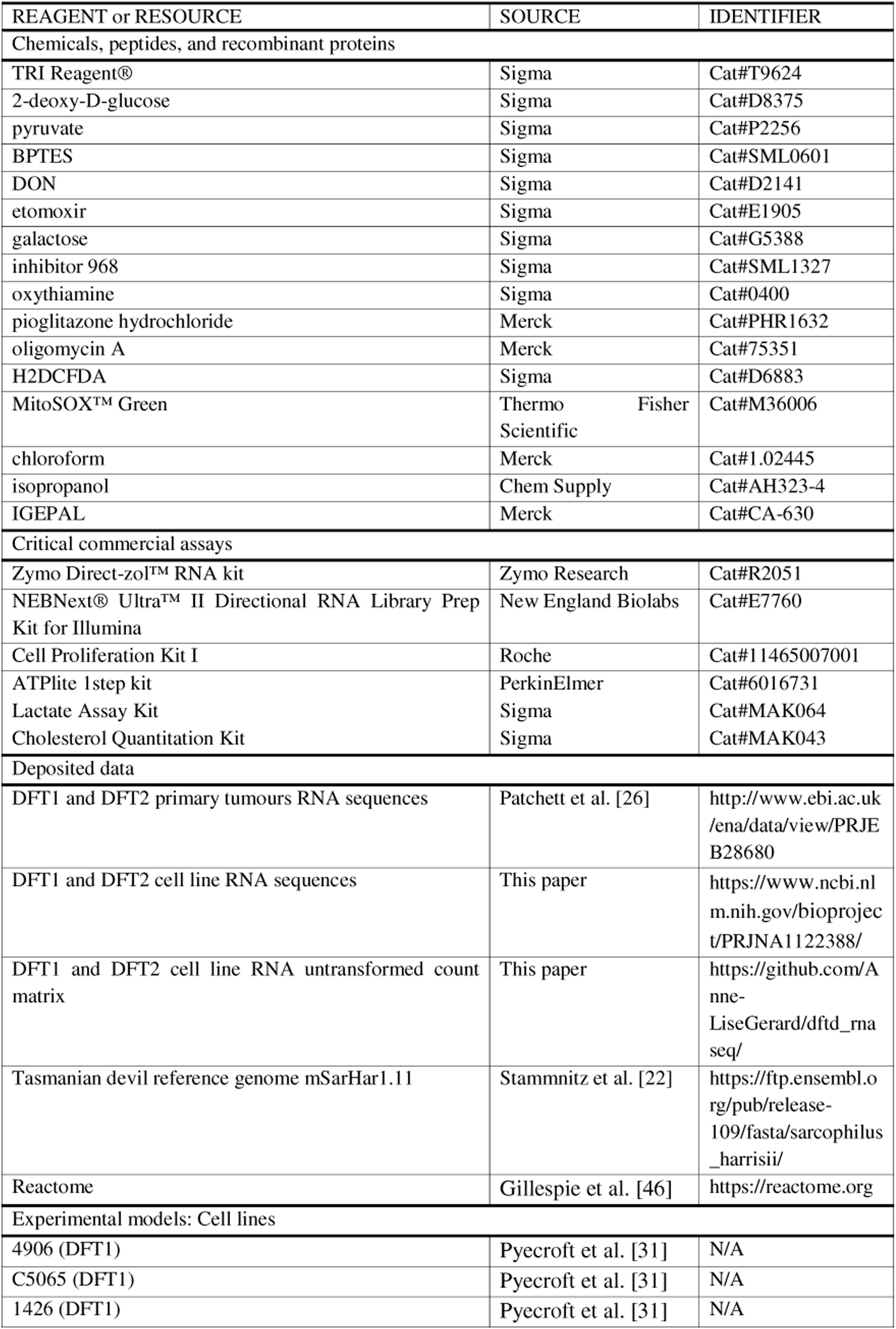
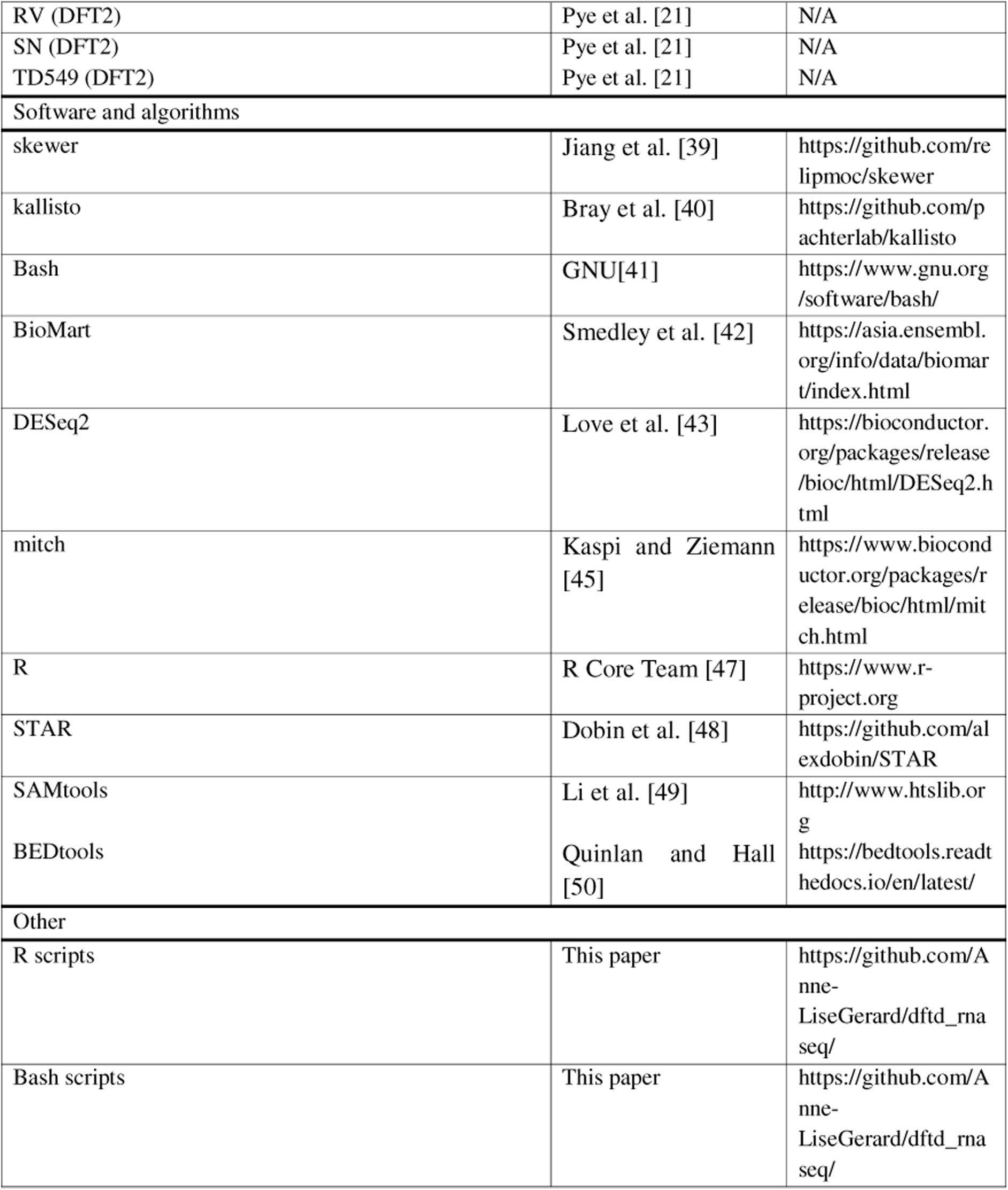
Key resources used in the study.

### QUANTIFICATION AND STATISTICAL ANALYSIS

#### Sequence analysis

Sequence reads were filtered (3’ end quality trimming with a quality threshold of 10) and trimmed using skewer [39] (v0.2.2) and pseudo-aligned to the Tasmanian devil reference genome (mSarHar 1.11) using kallisto [40] (v0.50.0), both using Bash [41] scripts. Technical replicates were combined and genes with a mean count of less than 10 across all samples were removed to exclude genes with extremely low expression. Human homologs of Tasmanian devil genes were obtained from Ensembl (release 109) using BioMart [42]. Differential gene expression analysis between DFT1 and DFT2 cell lines, and between DFT1 and DFT2 tumour biopsies was performed using DESeq2 [43] (v1.42.0). Differential expression p-values underwent false discovery rate (FDR) adjustment to account for multiple testing [44]. An adjusted p-value (FDR) < 0.05 was considered significant. To identify groups of genes (pathways) that are statistically over-expressed, a multi-contrast pathway enrichment analysis was performed using mitch [45] (v1.14.0) and Reactome [46] gene sets (V85) combined with two custom gene sets (data not shown). This allowed to directly compare differential gene expression between DFT1 and DFT2 cell lines to DFT1 and DFT2 biopsies. We also interrogated significant differentially expressed genes (DEGs) belonging to Reactome pathways investigated in our metabolic assays. Differential gene expression and pathway enrichment analyses were performed in R [47] (v4.3.3). For pathway enrichment analysis, an adjusted p-value (FDR) < 0.05 was considered significant.

#### Tumour purity calculation

To ensure tumour biopsies were not contaminated by host tissue, tumour purity was calculated and compared to that of the cell lines (which are clonal, hence not contaminated by host tissue). Tumour biopsy and cell line sequences were trimmed using skewer [39] then aligned to the mSarHar1.11 genome using the STAR aligner [48]. Files were converted from .sam to .bam, sorted by genomic location and indexed using SAMtools [49]. Mean coverage was then calculated over arbitrary 1kb non overlapping windows over the entire genome and over two large deletions proper to DFT1 and DFT2, using the BEDtools [50] coverage function. The DFT1 specific deletion is found at 3:192,161,000-216,866,000 (as per Stammnitz et al.[51]) and the DFT2 specific deletion at 5:141461000-175266000 (chosen from Stammnitz et al. [52]). Tumour purity was then calculated as (coverage deletion / overall coverage) x 100. Tumour purity calculations were performed using Bash [41] scripts and are available in Table S1.

#### Mixed effects models

All statistical analyses were performed in R [47] (version 4.3.0) using Gaussian mixed effects linear models. Response variables were either cell viability or metabolite production. To choose appropriate explanatory variables (cell lines, DFT), models were built following the Zuur protocol [53] (Table S3). Briefly, models with or without experimental replicate as a random intercept were built to account for possible batch effects between experiments. The most parsimonious fixed effects model was then selected. Models with the lowest Akaike’s Information Criterion (AIC) and highest AIC weights were selected. Model assumptions (i.e., homogeneity of variances and normality of residuals) were verified using diagnostic plots. Model selection showed all experiments required cell lines to be considered separately and could not be pooled by DFT due to variability in the results among cell lines of the same DFT. Model means and 95 % confidence intervals are reported in Table S4. Contrasts were performed using Tukey’s post hoc test, with Bonferroni correction for multiple testing (Table S4).

## ABBREVIATIONS

2DG: 2-deoxy-D-glucose
2DG6P: 2-deoxyglucose-6-phosphate
ADO: 2-aminoethanethiol dioxygenase
ACSL1/3/4: acyl-CoA synthetase long-chain family member 1/3/4
AIC: Akaike’s Information Criterion
AKT: protein kinase B
ATP: adenosine triphosphate
BTN: bivalve transmissible neoplasia
CO□: carbon dioxide
CPT1A: carnitine palmitoyltransferase 1A
dNTP: deoxyribonucleotide triphosphate
DEG(s): differentially expressed gene(s)
DFT: Devil Facial Tumour
DFT1: Devil Facial Tumour 1
DFT2: Devil Facial Tumour 2
DFTD: Devil Facial Tumour Disease(s)
DNA: deoxyribonucleic acid
ETC: electron transport chain
FBS: fetal bovine serum
FDR: false discovery rate
FSHR: follicle-stimulating hormone receptor
g: relative centrifugal force (×g)
GSH: glutathione
H2DCFDA: 2′,7′-dichlorodihydrofluorescein diacetate
HK3: hexokinase 3
HKDC1: hexokinase domain containing 1
kb: kilobase
MDS: multidimensional scaling
MHC: major histocompatibility complex
mRNA: messenger ribonucleic acid
MTT: 3-(4,5-dimethylthiazol-2-yl)-2,5-diphenyltetrazolium bromide
NADPH: nicotinamide adenine dinucleotide phosphate (reduced form)
OXPHOS: oxidative phosphorylation
PCR: polymerase chain reaction
PDGF: platelet-derived growth factor
PDGFRA: platelet-derived growth factor receptor alpha
PFKM: phosphofructokinase, muscle-type
PI3K: phosphoinositide 3-kinase
PPP: pentose phosphate pathway
RFU: relative fluorescence units
RIN: RNA Integrity Number
RNA: ribonucleic acid
ROS: reactive oxygen species
RPM: revolutions per minute
TCA: tricarboxylic acid (cycle)
µM / mM: micromolar / millimolar

## AUTHOR CONTRIBUTIONS

Conceptualization: A.-L.G., F.P., A.G.S., M.M., M.Z., B.U.

Data Curation: A.-L.G., F.P., C.V.

Formal Analysis: A.-L.G., F.P., C.V., A.M.D.

Funding acquisition: A.-L.G., M.M.,R.H., F.T., B.U.

Investigation: A.-L.G., F.P., C.V. Methodology: A.-L.G., F.P., M.M., M.Z.

Project administration: A.-L.G., U.B. Resources: A.-L.G., F.P., C.V., A.G.S., H.S., M.M., M.Z.

Software: A.-L.G., F.P., C.V., M.Z. Supervision: A.-L.G., F.T., B.U. Validation: A.-L.G., F.P., C.V. Writing – Original draft: A.-L.G. Writing – Reviewing & Editing. A.-L.G., F.P., C.V., A.M.D., A.G.S., R.H., H.S., F.T., M.M., M.Z., B.U.

## ACKNOWLEDGEMENTS

This work was supported by an ARC Linkage (LP170101105), ARC Decra (DE170101116), ARC DP (DP230100162), an ANR TRANSCAN (ANR--18--CE35--0009), the Save The Tasmanian Devil Eric Guiler Research Funds, Deakin University’s LES Blue Sky and SEBE Industry partnership funds, a CNRS International Research Project, a Morris Animal Foundation grant (D19ZO--413) and the Hoffmann Family. This research was supported by use of the Nectar Research Cloud, a collaborative Australian research platform supported by the NCRIS-funded Australian Research Data Commons (ARDC). The authors gratefully acknowledge the contribution to this work of the Victorian Operational Infrastructure Support Program received by the Burnet Institute.

During the preparation of this work the authors used Apple Intelligence and ChatGPT 5.2 to improve the readability and language of some sentences in the manuscript. After using these tools, the authors reviewed and edited the content as needed and take full responsibility for the content of the published article.

The graphical abstract was made using BioRender.

## DATA AVAILABILITY STATEMENT

All data and code are publicly available as of the date of publication. All original code has been deposited on GitHub and is available at https://github.com/Anne-LiseGerard as of the date of publication. RNA-seq data have been deposited at NCBI’s Sequence Read Archive (SRA) and are available under PRJNA1122388. The raw count matrix generated from this data is available at https://github.com/Anne-LiseGerard. This paper also analyses existing RNA sequences, available under PRJEB28680 on the SRA. Metabolic assay data are available upon request.

## SUPPORTING INFORMATION

**Figure S1.**
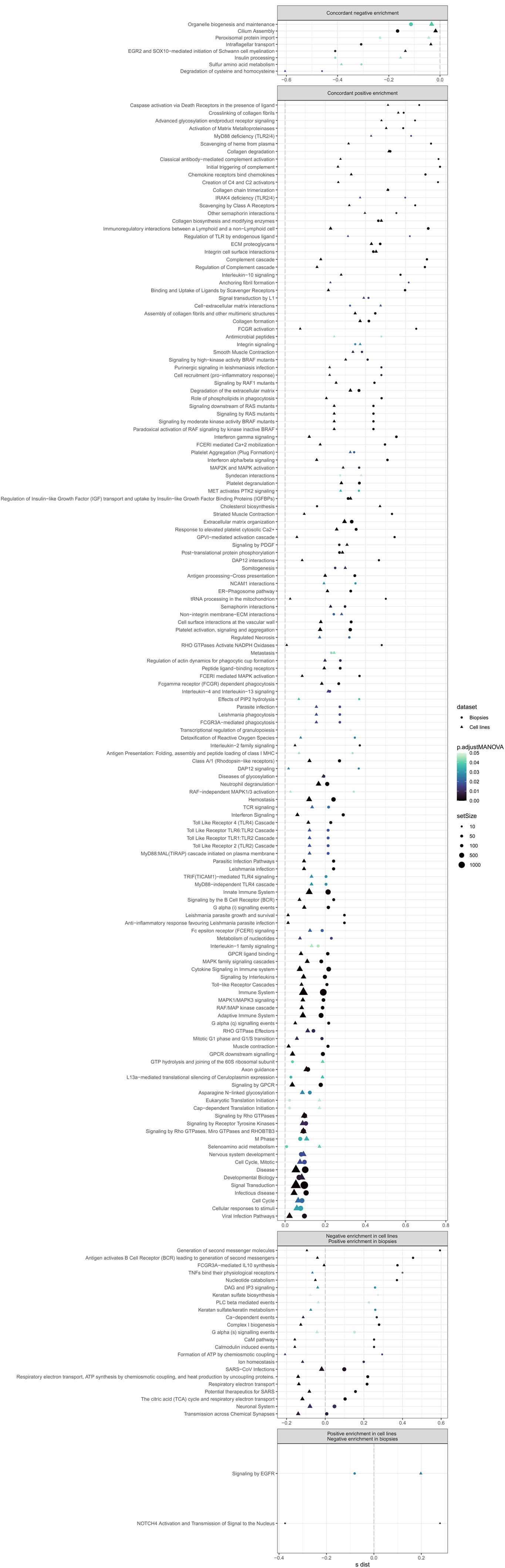
Multi-contrast pathway enrichment analysis using mitch.

Table S1. Tumour purity results.

Table S2. Gene list summary.

Table S3. Mixed effects model selection.

Table S4. Mixed effects models outputs.

