## Supplemental Figure 1 for "Metabolic pathways fuelling Devil Facial Tumour Disease"

SUPPLEMENTAL INFORMATION

Figure S1. Multi-contrast pathway enrichment analysis using mitch.
Multi-contrast pathway analysis performed with mitch on Reactome pathways, filtered for sets with an adjusted pvalue $< 0.05$. Each circle or triangle represents one gene set and is plotted according to its enrichment score (s-dist). Triangle or circle size is scaled according to the number of genes in a pathway (set size).

Table S1. Tumour purity results.
Devil Facial Tumour Disease cell line and tumour biopsy purity calculated based on large deletions proper to Devil Facial Tumour 1 (DFT1) and DFT2.

Table S2. Gene list summary.
Table containing all symbols, names and a brief description for genes mentioned in this article.

Table S3. Mixed effects model selection.
Table including degrees of freedom, Akaike's Information Criterion (AIC) and AIC weights, and which model was selected at each step.

Table S4. Mixed effects models outputs.
Table including estimates, lower confidence levels, upper confidence levels, and compact letter display of significant differences ($p < 0.05$). Significant differences were calculated using Tukey's post hoc test, with Bonferroni correction for multiple testing.

FigS1,

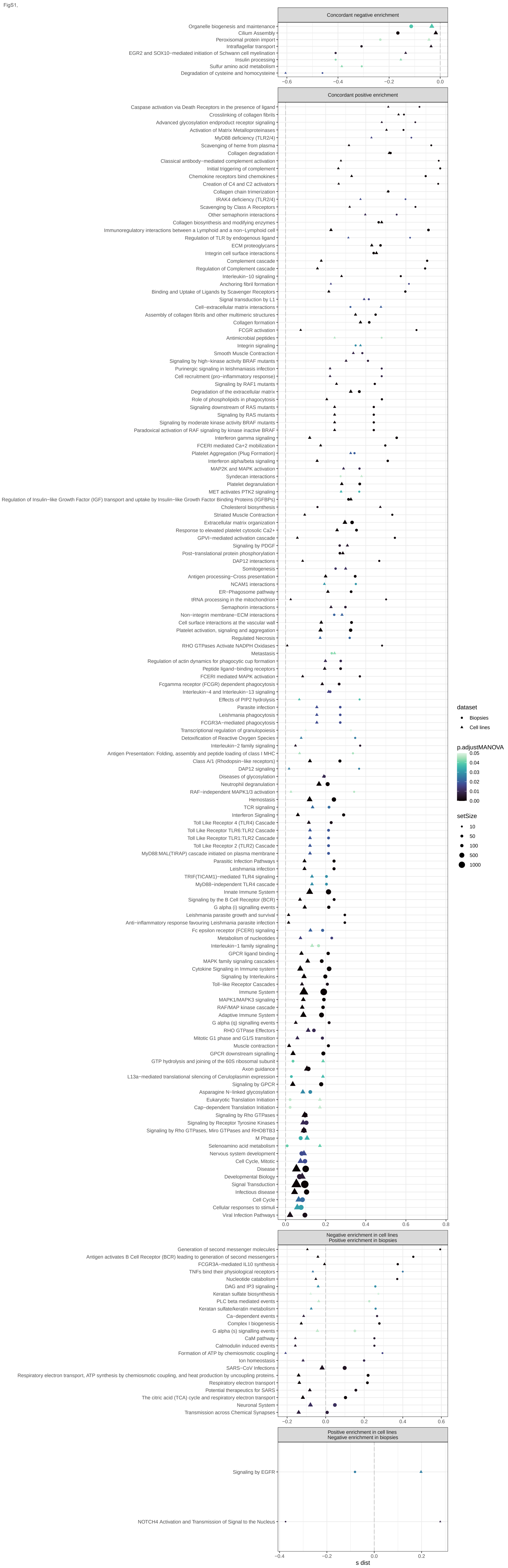
Concordant negative enrichment
Organelle biogenesis and maintenance
Cilium Assembly
Peroxisomal protein import
Intraflagellar transport
EGR2 and SOX10−mediated initiation of Schwann cell myelination
Insulin processing
Sulfur amino acid metabolism
Degradation of cysteine and homocysteine
−0.6
−0.4
−0.2
0.0
Concordant positive enrichment
Caspase activation via Death Receptors in the presence of ligand
Crosslinking of collagen fibrils
Advanced glycosylation endproduct receptor signaling
Activation of Matrix Metalloproteinases
MyD88 deficiency (TLR2/4)
Scavenging of heme from plasma
Collagen degradation
Classical antibody−mediated complement activation
Initial triggering of complement
Chemokine receptors bind chemokines
Creation of C4 and C2 activators
Collagen chain trimerization
IRAK4 deficiency (TLR2/4)
Scavenging by Class A Receptors
Other semaphorin interactions
Collagen biosynthesis and modifying enzymes
Immunoregulatory interactions between a Lymphoid and a non−Lymphoid cell
Regulation of TLR by endogenous ligand
ECM proteoglycans
Integrin cell surface interactions
Complement cascade
Regulation of Complement cascade
Interleukin−10 signaling
Anchoring fibril formation
Binding and Uptake of Ligands by Scavenger Receptors
Signal transduction by L1
Cell−extracellular matrix interactions
Assembly of collagen fibrils and other multimeric structures
Collagen formation
FCGR activation
Antimicrobial peptides
Integrin signaling
Smooth Muscle Contraction
Signaling by high−kinase activity BRAF mutants
Purinergic signaling in leishmaniasis infection
Cell recruitment (pro−inflammatory response)
Signaling by RAF1 mutants
Degradation of the extracellular matrix
Role of phospholipids in phagocytosis
Signaling downstream of RAS mutants
Signaling by RAS mutants
Signaling by moderate kinase activity BRAF mutants
Paradoxical activation of RAF signaling by kinase inactive BRAF
Interferon gamma signaling
FCERI mediated Ca+2 mobilization
Platelet Aggregation (Plug Formation)
Interferon alpha/beta signaling
MAP2K and MAPK activation
Syndecan interactions
Platelet degranulation
MET activates PTK2 signaling
Regulation of Insulin−like Growth Factor (IGF) transport and uptake by Insulin−like Growth Factor Binding Proteins (IGFBPs)
Cholesterol biosynthesis
Striated Muscle Contraction
Extracellular matrix organization
Response to elevated platelet cytosolic Ca2+
GPVI−mediated activation cascade
Signaling by PDGF
Post−translational protein phosphorylation
DAP12 interactions
Somitogenesis
Antigen processing−Cross presentation
NCAM1 interactions
ER−Phagosome pathway
tRNA processing in the mitochondrion
Semaphorin interactions
Non−integrin membrane−ECM interactions
Cell surface interactions at the vascular wall
Platelet activation, signaling and aggregation
Regulated Necrosis
RHO GTPases Activate NADPH Oxidases
Metastasis
Regulation of actin dynamics for phagocytic cup formation
Peptide ligand−binding receptors
FCERI mediated MAPK activation
Fcgamma receptor (FCGR) dependent phagocytosis
Interleukin−4 and Interleukin−13 signaling
Effects of PIP2 hydrolysis
Parasite infection
Leishmania phagocytosis
FCGR3A−mediated phagocytosis
Transcriptional regulation of granulopoiesis
Detoxification of Reactive Oxygen Species
Interleukin−2 family signaling
Antigen Presentation: Folding, assembly and peptide loading of class I MHC
Class A/1 (Rhodopsin−like receptors)
DAP12 signaling
Diseases of glycosylation
Neutrophil degranulation
RAF−independent MAPK1/3 activation
Hemostasis
TCR signaling
Interferon Signaling
Toll Like Receptor 4 (TLR4) Cascade
Toll Like Receptor TLR6:TLR2 Cascade
Toll Like Receptor TLR1:TLR2 Cascade
Toll Like Receptor 2 (TLR2) Cascade
MyD88:MAL(TIRAP) cascade initiated on plasma membrane
Parasitic Infection Pathways
Leishmania infection
TRIF(TICAM1)−mediated TLR4 signaling
MyD88−independent TLR4 cascade
Innate Immune System
Signaling by the B Cell Receptor (BCR)
G alpha (i) signalling events
Leishmania parasite growth and survival
Anti−inflammatory response favouring Leishmania parasite infection
Fc epsilon receptor (FCERI) signaling
Metabolism of nucleotides
Interleukin−1 family signaling
GPCR ligand binding
MAPK family signaling cascades
Cytokine Signaling in Immune system
Signaling by Interleukins
Toll−like Receptor Cascades
Immune System
MAPK1/MAPK3 signaling
RAF/MAP kinase cascade
Adaptive Immune System
G alpha (q) signalling events
RHO GTPase Effectors
Mitotic G1 phase and G1/S transition
Muscle contraction
GPCR downstream signalling
GTP hydrolysis and joining of the 60S ribosomal subunit
Axon guidance
L13a−mediated translational silencing of Ceruloplasmin expression
Signaling by GPCR
Asparagine N−linked glycosylation
Eukaryotic Translation Initiation
Cap−dependent Translation Initiation
Signaling by Rho GTPases
Signaling by Receptor Tyrosine Kinases
Signaling by Rho GTPases, Miro GTPases and RHOBTB3
M Phase
Selenoamino acid metabolism
Nervous system development
Cell Cycle, Mitotic
Disease
Developmental Biology
Signal Transduction
Infectious disease
Cell Cycle
Cellular responses to stimuli
Viral Infection Pathways
0.0
0.2
0.4
0.6
0.8
Negative enrichment in cell lines
Positive enrichment in biopsies
Generation of second messenger molecules
Antigen activates B Cell Receptor (BCR) leading to generation of second messengers
FCGR3A−mediated IL10 synthesis
TNFs bind their physiological receptors
Nucleotide catabolism
DAG and IP3 signaling
Keratan sulfate biosynthesis
PLC beta mediated events
Keratan sulfate/keratin metabolism
Ca−dependent events
Complex I biogenesis
G alpha (s) signalling events
CaM pathway
Calmodulin induced events
Formation of ATP by chemiosmotic coupling
Ion homeostasis
SARS−CoV Infections
Respiratory electron transport, ATP synthesis by chemiosmotic coupling, and heat production by uncoupling proteins.
Respiratory electron transport
Potential therapeutics for SARS
The citric acid (TCA) cycle and respiratory electron transport
Neuronal System
Transmission across Chemical Synapses
−0.2
0.0
0.2
0.4
0.6
Positive enrichment in cell lines
Negative enrichment in biopsies
Signaling by EGFR
NOTCH4 Activation and Transmission of Signal to the Nucleus
−0.4
−0.2
0.0
0.2
s dist
dataset
Biopsies
Cell lines
p.adjustMANOVA
0.05
0.04
0.03
0.02
0.01
0.00
setSize
10
50
100
500
1000
